# Semaphorin 3C exacerbates liver fibrosis

**DOI:** 10.1101/2021.07.29.454292

**Authors:** Francesca De Angelis Rigotti, Lena Wiedmann, Max Ole Hubert, Margherita Vacca, Sana S. Hasan, Iris Moll, Silvia Carvajal, Wladimiro Jiménez, Maja Starostecka, Adrian T Billeter, Beat Müller-Stich, Gretchen Wolff, Bilgen Ekim-Üstünel, Stephan Herzig, Carolin Mogler, Andreas Fischer, Juan Rodriguez-Vita

## Abstract

**Background & Aims:** Chronic liver disease is a growing epidemic leading to fibrosis and cirrhosis. TGF-β is the pivotal pro-fibrogenic cytokine which activates hepatic stellate cells (HSC), yet, other molecules can substantially modulate TGF-β signaling in the course of liver fibrosis. Expression of the axon guidance molecules Semaphorins (SEMAs), which signal through Plexins and Neuropilins (NRPs), have been associated with liver fibrosis in HBV-induced chronic hepatitis. This study aims at determining their function in the regulation of HSCs.

**Approach & Results:** We analyzed publicly available patient databases and liver biopsies. We employed transgenic mice where genes are deleted only in activated HSCs to perform ex vivo analysis and animal models. SEMA3C is the most enriched member of the Semaphorin family in liver samples from cirrhotic patients. Higher expression of SEMA3C in patients with NASH, alcoholic hepatitis or HBV-induced hepatitis discriminates those with a more pro-fibrotic transcriptomic profile. SEMA3C expression is also elevated in different mouse models of liver fibrosis and in isolated HSCs upon activation. In keeping with this, deletion of SEMA3C in activated HSCs reduces myofibroblast marker expression. Conversely, SEMA3C overexpression exacerbates TGF-β-mediated myofibroblast activation, as shown by increased SMAD2 phosphorylation and target gene expression. Among SEMA3C receptors, only NRP2 expression is maintained upon activation of isolated HSCs. Interestingly, lack of NRP2 in those cells reduces myofibroblast marker expression. Finally, deletion of either SEMA3C or NRP2, specifically in activated HSCs, reduces liver fibrosis in mice.

**Conclusion:** SEMA3C is a novel marker for activated HSCs that plays a fundamental role in the acquisition of the myofibroblastic phenotype and liver fibrosis.

## Introduction

Chronic liver disease (CLD) is a global epidemic that affects more than 850 million people(1). The cause of CLD can be attributed to different origins, most importantly hepatitis virus C and B, alcohol consumption and metabolic disorders leading to non-alcoholic steatohepatitis (NASH)(2). CLD is a complex disorder with different cell types contributing in multiple ways to its progression(2). However, no matter the cause for CLD, all etiologies have in common that hepatic stellate cells (HSCs) get activated. These produce aberrant extracellular matrix (ECM), accumulating in the extracellular space, generating a scar(3) and eventually impairing liver function(4). Understanding how HSCs get activated could help design better therapies to ameliorate the consequences of CLD.

It is well known that TGF-β is the main driver for the activation of HSCs(4). TGF-β signals through its receptor complex to activate mainly SMAD2/3 transcription factors(5). The TGF-β receptor complex is composed of the different TGF-β receptors, type I and II, and can include co-receptors like Neuropilin-1 (NRP1) or NRP2. NRPs exert no signaling function on their own, but modulate the signaling machinery(6–8). These co-receptors can also be recognized by other signaling molecules, such as distinct semaphorin proteins, suggesting that these may alter TGF-β signaling.

The semaphorin family of proteins (SEMA) were initially described to be involved in axon guidance during development(9), however, these proteins contribute to several other cell functions(10). The class III semaphorin (SEMA3) subfamily is the only group of semaphorins that is secreted(11). In chronic obstructive coronary disease, the expression of SEMA3F on the surface of neutrophils has been shown to retain neutrophils in the inflamed tissue, delaying the resolution of inflammation. On the contrary, when neutrophils express Semaphorin 4D on their surface their activation is reduced(12). This opposite functions in neutrophil activation suggests that each semaphoring member can have its own function and that the deeper knowledge of each member of this family could contribute to a better understanding of human disease. Some of SEMA3 members have been reported to be involved in the regulation of fibrosis. SEMA3A activates retinal fibroblasts, inducing differentiation into myofibroblast(13), while SEMA3E mediates HSC activation in a model of acute liver injury(14). Recently, SEMA3C expression has been shown to correlate with liver cirrhosis upon hepatitis B virus (HBV) infection(15). However, it has not suggested any functional role in this process. Interestingly, SEMA7A, a member of a different class of semaphorins, has been shown to potentiate TGF-β-induced liver and pulmonary fibrosis(16,17), suggesting that SEMAs could have a role in the regulation of TGF-β. Intriguingly, SEMA3C signals mainly through its interaction with PlexinD1, NRP1 and NRP2(18). While NRP1 plays an essential role in liver fibrosis(19,20), the role of NRP2 in liver fibrosis is unknown. Here we show that SEMA3C potentiates TGF-β signaling and exacerbates myofibroblast trans-differentiation induced by TGF-β and that this is, at least in part, mediated by NRP2.

## Methods

### Patient samples

Liver biopsies were collected from patients at the Klinikum rechts der Isar, Technical University Munich (TUM) Collection of the samples was approved by the tissue bank and the local ethical committee of the medical faculty (160/19S) in accordance with the Declaration of Helsinki. All patients gave preoperative consent. Samples collection include formalin fixed paraffin embedded tissue from patients with either steatosis (n=8), chronic viral hepatitis B (HBV; n=10) hepatitis C (HCV; n=12) or history of alcohol abuse (n=14).

### Mouse models

All animal experiments were approved by and performed in accordance with the institutional and regional Animal Welfare Committee. Animals were housed in groups in pathogen-free conditions. SM22α^CRE^/SEMA3C^fl/fl^ or NRP2^fl/fl^ constitutive knockout mice were obtained by crossing SM22α^CRE^ mice with either SEMA3C^fl/fl^ or NRP2^fl/fl^ mice to achieve the specific deletion of *Sema3C* or *Nrp2* genes only in cells expressing SM22α. These mice did not show any evident burden attributable to their genetic background at the ages at which the experiments were carried out. A constitutive knockout was chosen to fit to the continuous character of the disease, resulting in a knockout of SEMA3C or NRP2 in all activated HSCs independent of when their activation happened. Since SM22α is expressed in cardiomyocytes and both, SEMA3C and NRP2 play important roles in cardiac function and development (21,22), mice had to be euthanized before any potential burden could appear in the animals (around the 30 weeks of age) according to Animal Welfare Committee regulations. This decision was also based on the findings that 30 weeks old SM22α^CRE^/NRP1^fl/fl^ mice develop heart failure (23). For genotyping, the following primers were used: SM22α^CRE^ fw: GCGGTCTGGCAGTAAAAACTATC, rev: GTGAAACAGCATTGCTGTCACTT; SEMA3C flox fw: GAATCTGGCAAAGGACGATG, rev: GACCACTGGGCTTGAGAGAG; NRP2 flox common fw: AGCTTTTGCCTCAGGACCCA, NRP2 flox mutant rev: CCTGACTACTCCCAGTCATAG NRP2 Wildtype rev: CAGGTGACTGGGGATAGGGTA. Experiments started between 8-12 weeks of age on male mice. All mice were obtained from in-house breeding.

For CCl_4_ treatment, mice were kept on standard chow and injected 3x weekly with 0,5 mg/kg CCl_4_ (VWR International, Cat. No. SIAL289116) diluted 1:5 in olive oil (Carl Roth, Cat. No. 8873.1) for 2, 4 or 6 weeks.

For high-fat diet (HFD) experiments, wild type C57BL/6N mice were kept on a diet containing 60% fat, 20% carbohydrate, 20% proteins (Research Diets, Cat. No. D12492i) for 13 weeks from the age of 8 weeks. Control mice received standard chow.

For the FPC diet, wild type C57BL/6N mice were fed for 24 weeks with Fructose-Palmitate-Cholesterol high fat diet (Research Diets, Cat. No. D17020104i), while littermate controls were kept on control diet (Research Diets, Cat. No. D17020105i).

### Cells

GRX, a cell line of activated HSCs derived from a mouse model of schistosomiasis(24), and primary hepatic stellate cells (HSCs) were used in this work. Cells were cultured in DMEM GlutaMax 1mg/L glucose (Thermo Fisher Scientific, Cat. No. 21885025) supplemented with 10 % FCS and 1 % Pen/Strep. For the experiments with TGF-β, GRX cells were maintained in starvation (medium without FCS) for 48 h before stimulation and during all the duration of the experiment. Stimulation with TGF-β (10 ng/ml, Bio-Techne, Cat. No. 240-B-010/CF) was performed for 15 minutes, 3 or 24 hours.

### HSC isolation

Isolation of primary murine hepatic stellate cells (HSCs) was performed as previously described with individual modifications(25). In short: After cervical dislocation and immediate median laparotomy, the inferior vena cava (IVC) was exposed and then liver perfusion was performed via the IVC for 2 minutes with EGTA buffer(25), for 5 minutes with pronase (MERCK, cat. no. 10165921001, 14 mg in 35 ml of enzyme buffer solution (25) per mouse) and for 5 minutes with liberase (MERCK, cat. no. 5401127001, 1,25mg in 40 ml of enzyme buffer solution (25) per mouse). Afterwards, the liver capsule was carefully removed and the liver was minced. The minced liver was resuspended in GBBS/B(25) and then washed twice. HSCs were separated from other parenchymal and non-parenchymal liver cells by density gradient centrifugation, using Histodenz (MERCK, cat. no. D2158-100G, 4.94 g of Histodenz in 15 ml of GBSS/A(25)) for layering. HSCs were collected from the interphase between the two layers and then plated in 10% FCS, 1% PenStrep DMEM. For 0-9 days activation experiments, a fraction of HSCs was lysed and RNA was isolated immediately after HSC collection. Remaining HSCs were cultured for 9 days and then lysed for RNA isolation. Before to be used in experiments, HSC cells that were isolated from SM22α^CRE^/SEMA3C^fl/fl^ mice were tested for gene recombination by PCR. The following primers were used: AAGGCGCATAACGATACCAC and GACCACTGGGCTTGAGAGAG. Recombined gene gave a band around 250bp, while the absence of recombination gave a band around 1000bp.

### Lentiviral overexpression

Sema3C cDNA (NCBI DQ890847) vector from Genomic and Proteomics Core Facility of German Cancer Research Center was shuttled into pLenti6.2/V5-DEST (Invitrogen) by Gateway cloning for virus production. Sema3C or empty plasmids were transfected with plasmids for envelope (pMD2.G plasmid coding for VSV-G) and packaging (psPAX2) of lentiviral particles into HEK293T cells for virus production. Supernatant of these HEK293T cells was harvested 48 and 72h after transfection and virus was enriched by ultracentrifugation. Stable cell lines were generated by infection of GRX cells with viral particles and subsequent selection with blasticidin.

### Immunohistochemistry

Livers of mice were harvested and fixed in ROTI®Histofix 4 % (Carl Roth GmbH, Cat. No. P087.3) over night at 4°C before paraffin embedding. Stainings were performed on deparaffined cuts of 3µm.

Sirius Red staining was performed according to manufacturer’s protocol (Picro-Sirius Red Stain Kit, Dianova, Cat. No. PSR-1; Picro-Sirius Red Staining Kit, Morphisto, Cat. No. 1342500250). For SEMA3C (Bio-Techne, Cat. No. AF1728, 1:50), DESMIN (Abcam, Cat. No. ab15200, 1:100) and SM22α (abcam, Cat. No. ab137453, 1:100) stainings, antigen retrieval was performed at pH=6. After 3 fast washes in permeabilization buffer (TBS-0,025% Triton-X100), tissues were incubated for 30 minutes in Animal free Blocking solution (Cell Signaling 15019S, 1:5) and then with primary antibodies at 4°C ON. Endogenous peroxidase activity was inhibited by H_2_O_2_ treatment before adding Goat IgG VisUCyte HRP Polymer (R&D Systems, Cat. No. VC004 Substrate: DAB Substrate Kit, Zytomed, Cat. No. DAB057) for 30 minutes. For double staining, samples were subsequently incubated for 30 minutes with ZytoChem Plus (AP) Polymer anti-Rabbit (Biozol Diagnostica, Cat. No. ZYT-ZUC031-00, Substrate: Permanent AP-Red-Kit, Zytomed, Cat. No. ZUC001-125). Tissue was counterstained with hematoxylin for 3 seconds before washing, dehydration and embedding in mounting medium (Eukitt, Cat. No. 202200401). For immunofluorescence of αSMA, antigen retrieval was performed at pH=9. After 3 fast washes in permeabilization buffer (TBS-0,025% Triton-X100), samples were incubated for 30 minutes in 10% goat-serum, (Cell Signaling 15019S, 1:5) and then with diluted anti-αSMA-Cy3 antibody (Sigma Aldrich, Cat. No. C6198, 1:200) for 1h at RT. After washes, tissue was dehydrated and mounted with fluorescence mounting medium (Dako, Cat. No. S3023). Pictures were acquired using the Zeiss Axio Scan.Z1 and analyzed with the Fiji Software.

### Immunocytochemistry

For immunofluorescence staining of αSMA, GRX cells were seeded onto coverslips, starved ON and stimulated with 10 ng/ml TGF-β (Bio-Techne, Cat. No. 240-B-010/CF) for 24 h. Cells were fixed and permeabilized using cold MeOH for 3 min on ice. After blocking, cells were incubated with αSMA antibody (Sigma, Cat. No. A5228, 1:500) (**Suppl. Table 1**) for 2 hours at room temperature. Donkey anti-mouse IgG Alexa Fluor 546 antibody (Invitrogen, Cat. No. A10036) was incubated for 1 h prior to DAPI staining. Coverslips were mounted on slides with fluorescence mounting medium (Dako, Cat. No. S3023). Images were acquired using Zeiss Axio Scan.Z1.

### mRNA isolation and qRT-PCR

For qRT-PCR analysis, mRNA of GRX cells was isolated with the innuPREP RNA Mini Kit (Analytik Jena AG, Cat. No. 845-KS-20400250) according to manufacturer’s protocol. For HSCs, the ARCTURUS® PicoPure® RNA Isolation Kit (Thermo Fisher Scientific, Cat. No. KIT0214) was used. The extraction of mRNA from snap-frozen tissue was performed using Trizol® reagent (Thermo Fisher Scientific, Cat. No. 15596026) and precipitation with isopropanol. Retrotranscription of mRNA was carried out using the Reverse Transcriptase Kit (Applied Biosciences, Cat. No. 4368814). qRT-PCR was performed using SybrGreen Master Mix (Applied Biosciences, Cat. No. A25742) and primers listed in **Suppl. Table 2**. Fold changes were assessed by 2^−ΔΔ*C*t^ method and normalized with the gene *Cph* or *Hprt*.

### Western Blot

Western blot was performed as previously described. In short, cultured cells were lysed with Cell Lysis Buffer (Cell Signaling, Cat. NO. 9803S) supplemented with 2% PMSF (50mM in EtOH). For snap-frozen livers, tissue was homogenized in RIPA buffer (25mM Tris pH 7,4, 150mM NaCl, 1% NP40, 1% Sodium Deoxycholate, 0,1% SDS) supplemented with 2% PMSF (50mM in EtOH). Proteins separated via SDS-PAGE gels were blotted onto PVDF membranes (Millipore, Cat. No. ISEQ00010). To reduce non-specific antibody binding and reduce background, blocking-and antibody-incubations were performed in TBS + 0,1% Tween20 + 5% milk. Following primary antibodies (**Suppl. Table 1**) were used: anti-NRP1 (1:1000, R&D Systems, Cat. No. AF566), anti-NRP2 (1:1000, R&D Systems, Cat. No. AF567), anti-VCP (1:2000, abcam, Cat. No. ab109240), anti-pSMAD2/3 (1:1000, Cell Signaling, Cat. No. 8828), anti-SM22α (1:1000, abcam, Cat. No. ab137453), anti-FN1 (1:1000, abcam, Cat. No. ab2413). After washes, appropriate HRP-coupled secondary antibodies (rabbit anti-mouse, Cat. No. P0260, and goat anti-rabbit, Cat. No. P0448, from DAKO and rabbit anti-goat, Cat. No. HAF017, from R&D Systems) were used with a dilution 1:2500. HRP-conjugated rabbit anti-mouse (Dako, Cat. No. P0260) was also used to recognize heavy-chain immunoglobulins (hc-Ig) present in liver lysates. SuperSignal™ West Pico PLUS Chemiluminescent Substrate (Thermo Fisher Scientific, Cat. No. 34579) was used to detect HRP.

### Gene set enrichment analysis (GSEA)

Gene Set Enrichment Analysis (GSEA, Broad Institute)(26) was used to examine differentially expressed genes (DEGs). The output of GSEA is an enrichment plot (ES), a normalized enrichment score (NES) which accounts for the size of the gene set being tested, a p-value, and an estimated False Discovery rate (FDR). We computed p-values using 1,000 permutations for each gene set and corrected them with the false-discovery rate (FDR) method. When several probe sets were present for a gene, the mean of the probe set was used. The patient classification based on the expression of a gene was performed using the median ox expression as a threshold. Patients were classified as over or under the median value. We used publicly available data sets of liver fibrosis patients (NCBI GEO GSE45050, GSE49541, GSE61260, GSE103580, GSE83898) for the analysis.

### Statistical analysis

Data are presented as mean ± SD, with data points representing different biological replicates. Statistical analysis was performed with GraphPad Prism 9 software. The test used in each experiment is described in the figure legend. For *in vivo* experiments, the non-parametric Mann-Whitney test was applied, given the low number of samples used (8 or less). For *in vitro* experiments, Gaussian distribution of data was assumed, so parametric unpaired t-test was performed. For comparing *ex vivo* analysis of HSC activation comparing 0 and 9 days after isolation, paired t-test was carried out. One-tailed analysis was performed only when one direction hypothesis was supported by previously published data. Differences were considered significant, if p-value was ≤0.05. Asterisks were used as followed: * p ≤0.05, ** p ≤ 0.01, *** p ≤ 0.001.

## Results

### SEMA3C expression is associated with liver fibrosis in human patient samples

To address relative gene expression patterns of semaphorin family members in human CLD samples, we examined a publicly available database (GSE45050) which provides gene expression profiles from patients with liver cirrhosis (n=5) and healthy donors (n=3). Gene set enrichment analysis (GSEA) comparing healthy versus cirrhotic patients using the GO_SEMAPHORIN_PLEXIN_SIGNALING _PATHWAY gene set revealed that this gene set was enriched upon liver cirrhosis (**Suppl. Figure 1A**). The most enriched gene in this set was SEMA3C, suggesting that its role could be important in the context of liver cirrhosis. Surprisingly, SEMA7A, whose function had been suggested to potentiate liver fibrosis, was not among the enriched genes in this analysis. This analysis, as part of the process, performs a rank with all the genes expressed from most enriched in healthy to most enriched in cirrhotic. Using this rank, we extracted a list of the 500 most enriched genes in cirrhotic samples as a signature of cirrhosis and named it “Cirrhosis signature”. Interestingly, three different class 3 semaphorins were among these genes (**Suppl. Table 3**): SEMA3D, SEMA3A, among the top 400, and SEMA3C, which was in the top 100 of most enriched genes. However, this cohort is quite limited. The number of patients is rather low, and the etiology of the cirrhosis was not disclosed(27). Therefore, we decided to test whether our signature could identify fibrotic patients before patients were showing signs of cirrhosis, we employed a dataset of patients suffering NAFLD with different grades of fibrosis (GSE49541). We classified these patients in terms of their fibrotic grade in mild (0/1) or severe (3/4)(28). When performing GSEA using the Cirrhosis signature we observed a significant enrichment of this signature in the patients showing severe fibrosis (**Suppl. Figure 2B**), which demonstrates that this signature is a good tool to determine a fibrotic expression profile in patients with CLD. Importantly, when performing GSEA using the go_semaphorin_plexin_signaling _pathway gene set in this classification of patients we observed that it was enriched in those patients diagnosed with a severe fibrosis (**Figure 1A**). Moreover, SEMA3C was again the most enriched gene in this analysis (**Figure 1A**), further supporting the potential role of this gene in liver fibrosis development. It is worth noting that SEMA7A was again not present among the most enriched genes. We used other publicly available datasets from different cohorts of patients suffering different levels of NAFLD including NASH (GSE61260) (**Figure 1B**)(29), alcoholic hepatitis (GSE103580) (**Figure 1C**)(30) and chronic HBV infection (GSE83898) (**Figure 1D**)(31). We classified the different cohorts of patients based on their SEMA3C expression as over (SEMA3C-high) or under (SEMA3C-low) the median expression of SEMA3C among all patients. We removed SEMA3C from the cirrhosis signature gene set to avoid introducing a bias in our analysis. GSEA showed significant enrichment of this gene set in patients with higher SEMA3C expression in all etiologies for CLD (**Figure 1B-D**), suggesting its importance in liver fibrosis. To ensure that this was not an unspecific association between SEMA3C and fibrosis, we performed the same classification (over or under the median) with other semaphorin genes that have been related to fibrosis in the literature, SEMA3E, SEMA3A, SEMA5A and SEMA7A(13–16); we also included SEMA3D because it is present in the cirrhosis signature. We found that although some of these genes showed the same trend, SEMA3C had the strongest significant association in all three disorders (**Suppl. Figure 1C**). In summary, these data indicate that level of expression of SEMA3C is associated with a fibrotic mRNA expression profile in patients with CLD from different etiologies. To understand whether this was due to its impact on fibroblast activation, we examined the expression of SEMA3C in liver samples from patients suffering: non-alcoholic fatty liver disease (NAFLD), alcoholic cirrhosis, chronic HCV or chronic HBV infection. We performed a double staining with SEMA3C and α-smooth muscle actin antibodies and observed that they were often expressed in the same cells (**Figure 1E**), indicating that activated fibroblasts express SEMA3C. We further validated these results in other samples of NAFLD, where we also found SEMA3C expression in DESMIN positive cells (**Suppl. Figure 1D**), indicating that SEMA3C was expressed in HSCs. We also analyzed the expression of a marker for fibroblast activation, smooth muscle protein 22-alpha (SM22α). Although, the investigated patient samples exhibited only mild signs of fibrosis (based on clinical pathology reports), they started to show SM22α positive staining in non-arterial areas (**Suppl. Figure 1E**). This indicates that fibroblasts, probably HSCs, had been activated already.

**Figure 1.**
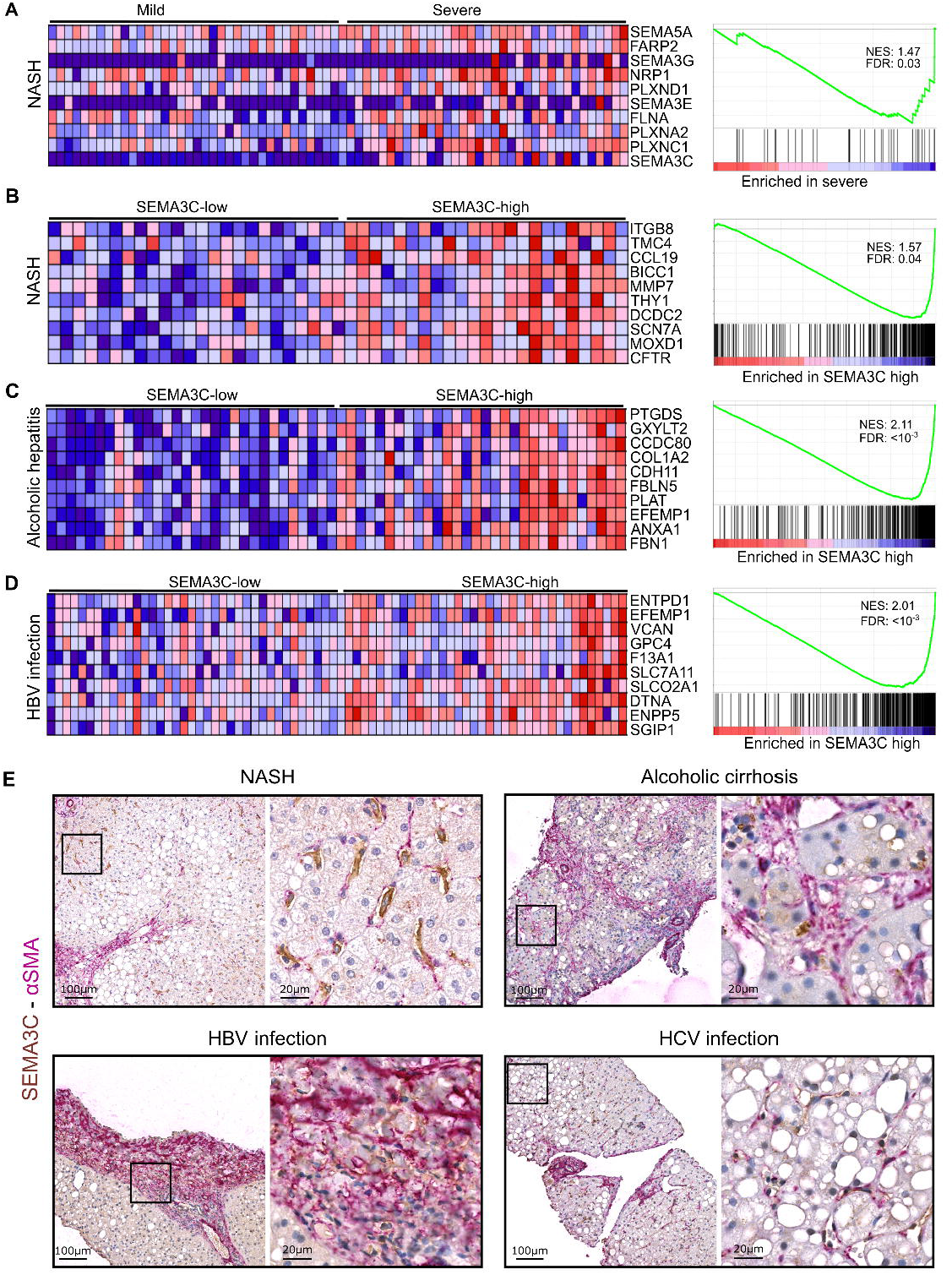
SEMA3C expression is associated with liver fibrosis in humans. **A** GSEA of the dataset GSE49541 using GO_SEMAPHORIN_PLEXIN_SIGNALING _PATHWAY gene set. Left panel shows a heatmap of the 10 genes most differentially enriched with the most enriched gene at the bottom. Right panel shows the enrichment plot including the Normalization Enrichment score (NES) and false discovery rate (FDR). **B-D** GSEA of the dataset GSE61260 (B) GSE103580 (C) GSE83898 (D) using “Cirrhosis signature” gene set obtained from A. Patients were classified according to their expression levels of SEMA3C as over (high) or under (low) the mean. Left panels show a heatmap of the 10 genes most differentially enriched with the most enriched gene at the bottom. Right panel shows the enrichment plots including the NES and FDR. **E** Immunohistochemistry for αSMA (stained by alkaline phosphatase AP) and SEMA3C (stained by DAB) were performed on sections of human livers from 4 different etiologies: NASH, alcoholic cirrhosis, HBV infection and HCV infection. Samples were counterstained with hematoxylin. Pictures show both proteins in perisinusoidal areas and more fibrotic areas. Scale bars are 100µm in the zoom-out images and 20 µm in the zoom-in images.

### SEMA3C is upregulated in several mouse models for liver fibrosis

We further analyzed whether we could observe the same kind of association in mouse models for liver fibrosis. We used two models for NAFLD, i.) high fat diet (HFD) and ii.) fructose, palmitate, cholesterol and trans-fat diet (FPC), and in addition the classical model of CCl_4_-induced liver fibrosis. Although all three models showed signs of early fibrosis, only CCl_4_-treated mice had advanced liver fibrosis in relatively short time (**Suppl. Figure 2A-C**). Nonetheless, in all three models we observed a significant increase of SEMA3C upon liver injury (**Figure 2A**). Similar to what we observed in patients with NAFLD, SEMA3C was expressed by DESMIN positive fibroblasts and was more concentrated in fibrotic areas (**Figure 2B**).

**Figure 2.**
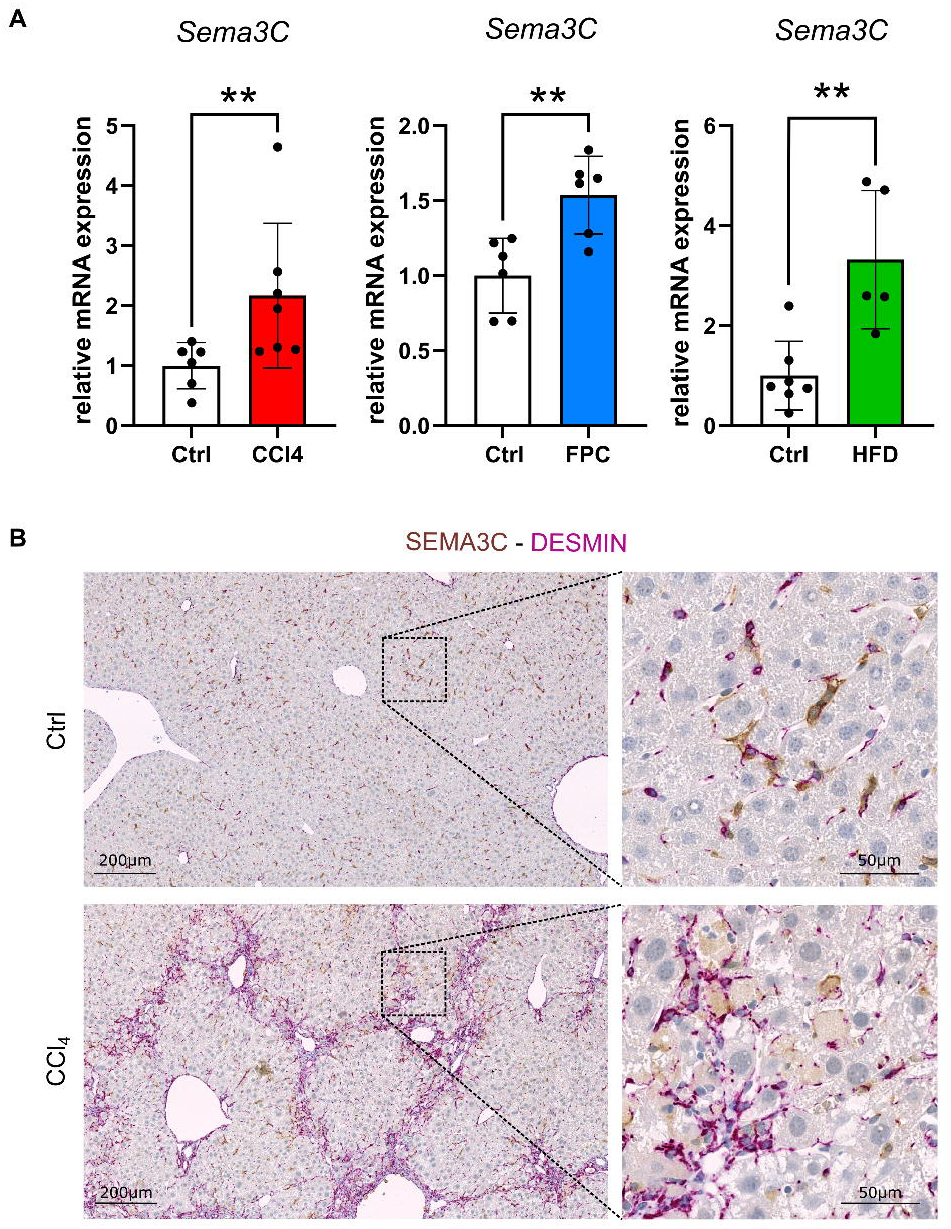
SEMA3C is upregulated during liver fibrosis in mice. **A** *Sema3C* expression was analyzed by qRT-PCR in 3 different models of liver fibrosis: 6 weeks of carbon tetrachloride-treatment (CCl_4_, column in red), 24 weeks of fructose-palmitate-cholesterol high fat diet (FPC, column in blue) and 13 weeks of high fat diet (HFD, column in green). Each mouse is denoted as a black dot and the mean ± SD of each group is represented as a column. White columns represent control mice. Two-tailed non-parametric Mann-Whitney test was performed to evaluate data significance. ** indicates p value <0,01. **B** DAB-SEMA3C staining (in brown) and AP-DESMIN staining (in purple) were performed on control livers and livers with advanced fibrosis (6 weeks CCl_4_-treatment). Tissue was counterstained with hematoxylin. Scale bars are 200 µm in the zoom-out images and 50µm in the zoom-in images.

### SEMA3C is a marker for HSC activation

Thus far, we have been able to correlate SEMA3C expression with liver fibrosis, but we aimed at understanding whether this was functionally relevant. For this reason, HSCs from livers of wild-type C57BL/6 mice were isolated by density gradient and activated by *in vitro* culture (**Suppl. Figure 3A**). *In vitro* culture has been shown to be sufficient to activate HSCs into myofibroblasts(32). We confirmed this by observing that several markers for activated fibroblasts (*Ctgf, Acta2* (αSMA), *Tagln* (SM22α) and *Pai-1)* were increased upon *in vitro* culture for 9 days (**Suppl. Figure 3B**). When looking at SEMA3C expression we observed a significant increase in activated HSCs (**Figure 3A**), indicating that SEMA3C can be considered as a new marker for activated fibroblasts.

**Figure 3.**
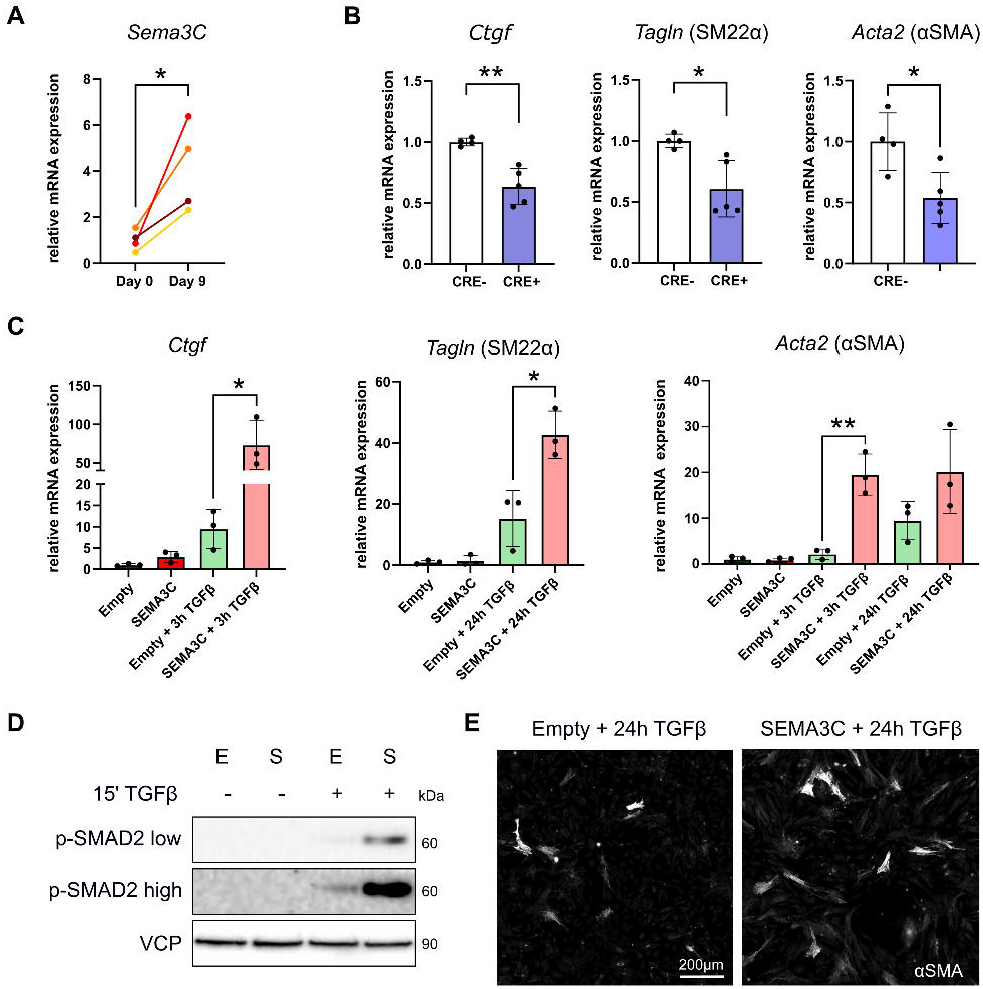
SEMA3C is a marker of hepatic stellate cells (HSC) and exacerbates fibroblast activation. **A** HSCs were isolated from WT mice and one portion was immediately frozen, while the other one was maintained in culture with complete medium for 9 days. Gene expression of *Sema3C* was analyzed in these samples by qRT-PCR. Two-tailed paired t-test was performed to evaluate data significance. **B** HSCs were isolated from SM22α^CRE^/SEMA3C^fl/fl^ mice and the control CRE-littermates, and cultured for 9 days. Gene expression of *Ctgf, Tagln* (SM22α) and *Acta2* (αSMA) was analyzed by qRT-PCR. Two-tailed unpaired t-test was performed to evaluate data significance. **C** SEMA3C-overexpressing GRX (SEMA3C, S) and control GRX (empty, E) were stimulated with TGF-β for 3 or 24 hours and gene expression of activation markers was evaluated by qRT-PCR. Two-tailed unpaired t-test was used to evaluate data significance. **D** Phosphorylation of SMAD2/3 was evaluated by Western blot, after 15 minutes of activation and VCP was used as loading control. Image is representative of 3 independent experiments. **E** Immunofluorescence of αSMA was performed on 24h-activated GRX. Scale bar is 200 µm. * and ** correspond to p value <0.05 and <0.01, respectively.

To demonstrate the importance of SEMA3C in HSC activation, we isolated HSCs from SM22α^CRE^/SEMA3C^fl/fl^ and their littermate controls SEMA3C^fl/fl^. In SM22α^CRE^/SEMA3C^fl/fl^ mice SEMA3C deletion is controlled by SM22α promoter, whose activity increases upon HSC activation. We observed that, after 9 days of culture, HSCs from CRE-positive mice had a significantly reduced expression of several myofibroblast markers compared to their CRE-negative littermate controls (**Figure 3B**), indicating that SEMA3C expression is necessary for HSC activation.

Further, we overexpressed SEMA3C in GRX cells, a cell line of activated HSCs(24). We observed that TGF-β-mediated gene expression was exacerbated in those cells overexpressing SEMA3C (**Figure 3C**). In addition, TGF-β-induced SMAD2 phosphorylation in SEMA3C-overexpressing GRX was significantly increased compared to control cells (**Figure 3D**). This was further confirmed by staining for αSMA by immunofluorescence. SEMA3C-overexpressing GRX showed a stronger formation of αSMA^+^ cytoskeletal filaments compared to control cells, which confirms the increased myofibroblast activation of HSCs upon SEMA3C overexpression (**Figure 3E**). All these data indicate that SEMA3C increases TGF-β responses in HSCs and GRX *in vitro*.

### SEMA3C exacerbates liver fibrosis in mice

To analyses whether SEMA3C could also mediate fibrosis *in vivo*, we induced liver fibrosis in mice lacking SEMA3C in myofibroblasts. For this, we employed SM22α^CRE^/SEMA3C^fl/fl^ mice. In CRE-positive mice, fibroblasts and all HSCs, which get activated at any phase of liver fibrosis progression, lack SEMA3C (**Figure 4A, Suppl. Figure 3A**). We used CCl_4_-injection as liver fibrosis model because, due to the short time this model requires to induce fibrosis, is the most compatible with the chosen mouse strain. Lack of SEMA3C in myofibroblasts reduced significantly collagen deposition, as shown by Sirius red staining of the livers (**Figure 4C**). In addition, mice those mice also showed a significant decrease of several liver fibrosis markers, such as *Acta2* mRNA expression, as well as FN1, SM22α and αSMA protein expression (**Figure 4B, D and E**). Moreover, immunoglobulin (hc-Ig) presence in the liver, which has been reported to be a marker for liver fibrosis(33) also in our CCL_4_ model (**Suppl. Figure 2C**), was strongly reduced (**Figure 4E**). Taken together, these data demonstrate that myofibroblast expression of SEMA3C exacerbates liver fibrosis.

**Figure 4.**
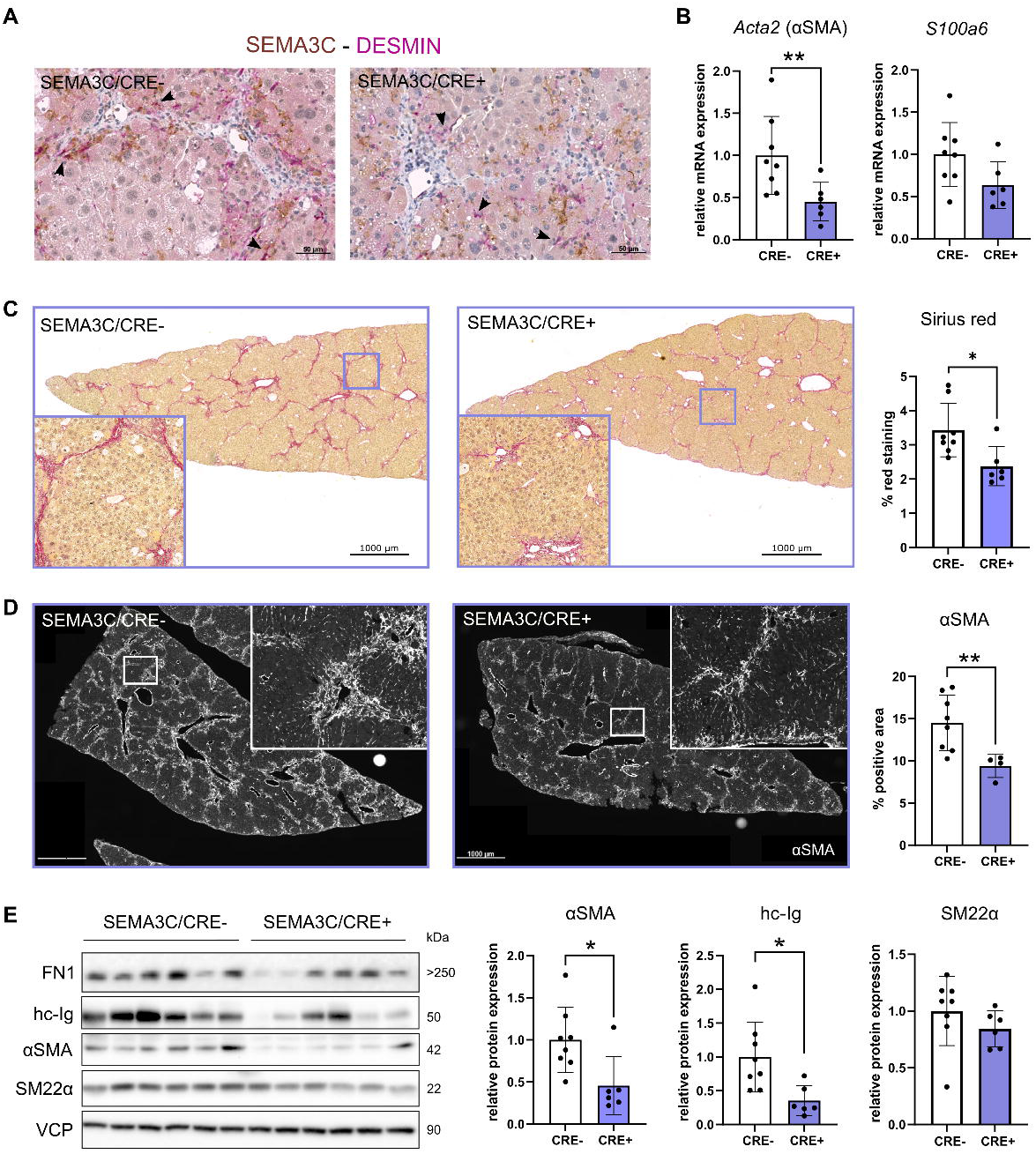
SEMA3C exacerbates liver fibrosis in mice. Liver fibrosis was induced in SM22α^CRE^/SEMA3C^fl/fl^ mice and their control CRE-littermates, by CCl_4_-treatment for 6 weeks. **A** SEMA3C staining (by DAB in brown) and DESMIN staining (by AP in purple) were performed to determine fibroblast-specific SEMA3C deletion (arrows) in CRE+ mice. Scale bars are 50µm. **B** Gene expression of *Acta2* (αSMA) and *S100a6* was analyzed in these samples by qRT-PCR. Liver fibrosis was also evaluated by Sirius red staining and immunofluorescence of αSMA. **C** Representative images and quantification of Sirius red staining are here shown, as well as representative images and quantification of αSMA staining (**D**). **E** The same samples were analyzed by Western blot. Fibronectin (FN1), heavy-chain immunoglobulin (hc-Ig), αSMA, SM22α and the loading control VCP are here shown and the bands quantified. Two-tailed non-parametric Mann-Whitney test was performed to evaluate data significance in the experiments of this figure. * and ** correspond to p value <0.05 and <0.01, respectively.

### NRP2 deletion in HSCs reduces liver fibrosis in mice

SEMA3C signals through PlexinD1, NRP1 and NRP2 receptors(34). Given that NRPs are co-receptors in the TGF-β receptor complex(35) we further determined their contribution to HSC activation and liver fibrosis. When analyzing the expression of NRPs upon fibrosis, we observed that NRP1 increased with fibrosis development, as expected from previous literature(19). Interestingly, we observed a similar upregulation of NRP2, suggesting a previously unknown effect of NRP2 on liver fibrosis (**Figure 5A**). However, when analyzing gene expression in primary HSCs activated in culture, we observed that while *Plxnd1* and *Nrp1* were almost abolished in activated HSCs, these conserved *Nrp2* expression (**Figure 5B**). These data suggest that NRP2 could be transducing the signal of SEMA3C in activated HSCs. To evaluate the relevance of NRP2 in HSC activation, we isolated HSCs from SM22α^CRE^/NRP2^fl/fl^ and their littermate controls NRP2^fl/fl^. We found several markers for HSC activation significantly reduced in cells lacking NRP2, in particular *Tagln* (SM22α) and *Acta2* (αSMA) (**Figure 5C**). Thus, we concluded that NRP2 is necessary for HSC activation *in vitro*.

**Figure 5.**
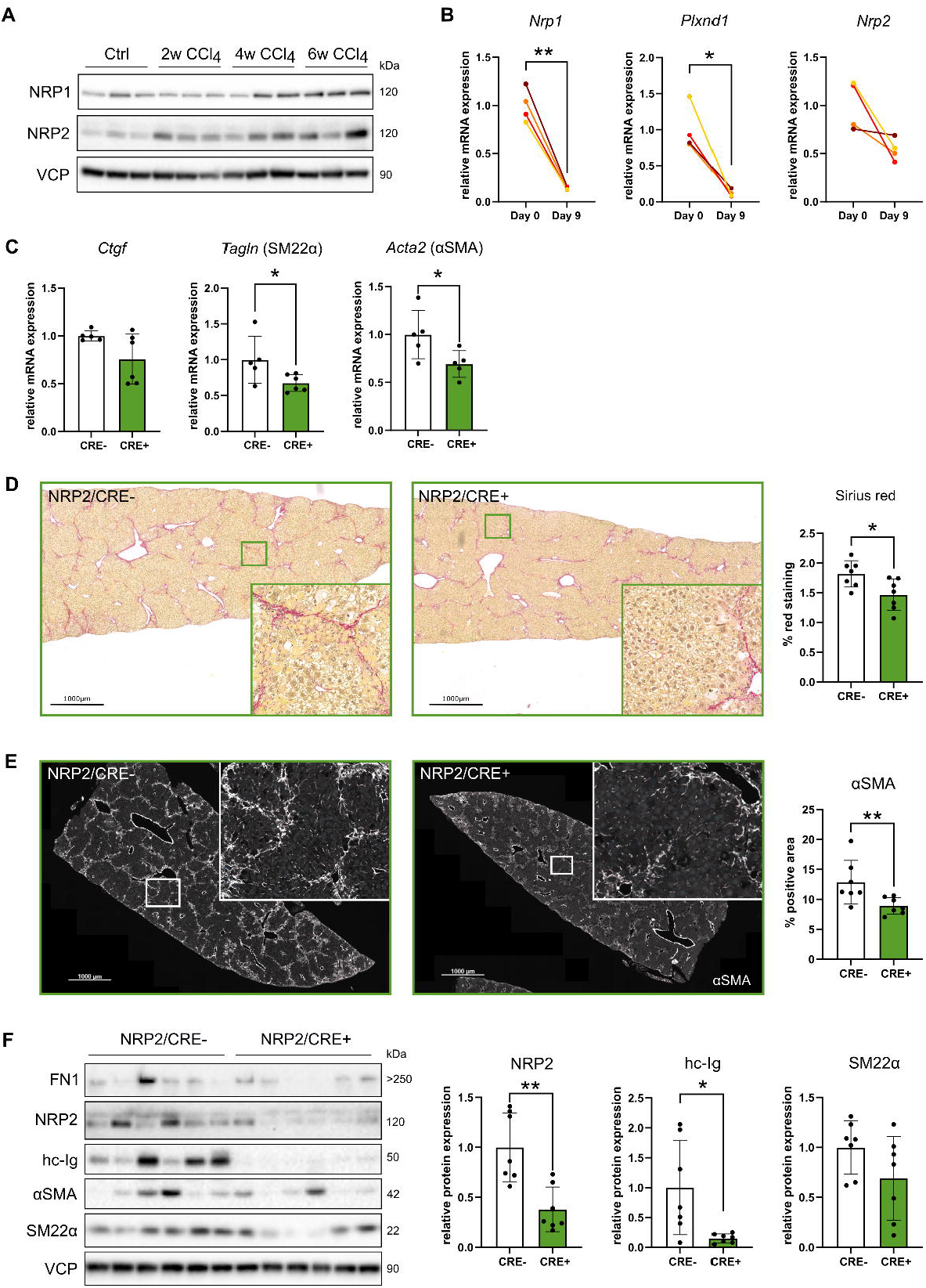
NRP2 on activated fibroblasts mediates fibrosis in mice. **A** Western blot was performed on whole liver tissues of mice treated for 2, 4 and 6 weeks (w) with CCl_4_ and their control littermates. Blots of Neuropilin-1 (NRP1), Neuropilin-2 (NRP2) and VCP are shown. **B** Gene expression of *Nrp1, Plxnd1* and *Nrp2* was analyzed by qRT-PCR in freshly-isolated WT-HSCs and after 9 days of culture. Two-tailed paired t-test was performed to evaluate data significance. **C** HSCs from SM22α^CRE^/NRP2^fl/fl^ mice and the control CRE-slittermates were cultured for 9 days and gene expression of *Ctgf, Tagln* (SM22α) and *Acta2* (αSMA) was examined by qRT-PCR. Two-tailed unpaired t-test was performed to evaluate data significance. **D** Liver fibrosis was induced in SM22α^CRE^/NRP2^fl/fl^ mice and the control CRE-littermates by CCl_4_-treatment for 6 weeks. Liver fibrosis was evaluated by Sirius red staining. Representative images of livers and quantification of the red staining by Fiji software are displayed for each group. **E** Liver fibrosis was also analyzed by immunofluorescence of αSMA and results are shown as representative images of livers and quantification of αSMA staining for the two groups. D and E panels, scale bars are 1000 µm. **F** The same samples were also investigated by Western blot. FN1, hc-Ig, αSMA and SM22α were used as markers for liver fibrosis and NRP2 was detected to verify knock-out of NRP2. Bands were quantified and normalized to VCP (graph on the right of the panel). Two-tailed non-parametric Mann-Whitney test was performed to evaluate the significance of *in vivo* data (panel D, E and F). * and ** correspond to p value <0.05, <0.01 respectively.

In order to confirm the important role of the co-receptor NRP2 in liver fibrosis, we injected CCl_4_ in mice lacking NRP2 only in myofibroblasts (SM22α^CRE^/NRP2^fl/fl^ mice) and their littermate controls to induce liver fibrosis. Lack of NRP2 in myofibroblasts reduced significantly collagen deposition, as shown by Sirius red staining of the livers (**Figure 5D**). We also observed a lower presence of several liver fibrosis markers in SM22α^CRE^/NRP2^fl/fl^ mice, such as SM22α, αSMA and immunoglobulin (**Figure 5E-F**). Therefore, the data suggest that the absence of NRP2 in activated HSCs reduces liver fibrosis.

## Discussion

CLD is a multifactorial disorder. Most of the molecular mechanisms underlying liver fibrosis are common among the different liver injuries triggering CLD. Also, most of them start with the transdifferentiation of HSCs into myofibroblasts. Understanding the mechanisms for HSC activation could lead to the discovery of better ways to diagnose and treat patients with CLD. Here we have uncovered a new marker for HSC activation, SEMA3C, which predicts a more advanced stage of the disease. In addition, we demonstrate that SEMA3C has a function in liver fibrosis progression, thereby constituting a potential therapeutic target itself. Moreover, we suggest that its effects are, at least in large parts, mediated by the receptor NRP2.

The so-called Western diet and sedentary lifestyle has enormously increased the rate of people affected by NAFLD, or more accurately described by metabolic dysfunction-associated fatty liver disease (MAFLD) (36). In addition, excessive alcohol consumption or viral infections contribute to the high rate of CLD, constituting a growing epidemic affecting almost a billion people worldwide(1). The discovery of safe treatments for CLD could be extremely beneficial. Here we describe a previously unknown role for SEMA3C in the activation of HSCs. We provide data showing that SEMA3C expression precedes other more obvious features of liver fibrosis. This suggests that SEMA3C expression might be employed for early detection of liver fibrosis and for a better classification of patients with risk of developing a more advanced disease before the deterioration of liver function arrives to a non-return point. Although liver biopsies are invasive, there is already a study suggesting that serum levels of SEMA3C associate with fibrosis in HVB infection(15), which implies that serum levels of SEMA3C could be employed for diagnosis in all other etiologies. Our data gives a mechanistic role for this potential diagnosis.

TGF-β is one of the main drivers for liver fibrosis. It is an important activator of HSCs, and its inhibition has been shown to block liver fibrosis progression(37). Unfortunately, TGF-β inhibitors are often too toxic to be administered due to pleiotropic effects in many organs. We found that SEMA3C exacerbates TGF-β responses in HSCs. We show how the overexpression of SEMA3C in GRX, a myofibroblast cell line derived from HSCs, increases TGF-β-induced gene expression and SMAD2 phosphorylation. Therefore, early increase of SEMA3C expression in HSCs could provoke an overreaction to TGF-β present in the extracellular compartment, accelerating the development of fibrosis that TGF-β mediates. Conversely, deletion of SEMA3C in HSCs reduced their level of activation *in vitro*. This indicates that SEMA3C is not only a consequence of HSC activation, it is also a driver for their activation itself. Future strategies aiming at blocking SEMA3C activity could be useful to reduce TGF-β responses only in damaged organs, limiting the toxicity by avoiding the inhibition of TGF-β pleiotropic actions. SEMA3C plays an important role in angiogenesis by inhibiting VEGF responses in endothelial cells (ECs) through NRP1(38). For this reason, it is important to separate the effects of SEMA3C on ECs from potential effects that SEMA3C could have in fibroblasts. To achieve this, we decided to use SM22α as a driver for CRE recombinase expression because it allowed us to restrict SEMA3C deletion to activated HSCs and myofibroblasts in general. This promoter is also active in VSMCs, although VSMC do not express SEMA3C (**Figure 2B**). Therefore, it is reasonable to suggest that our transgenic mice would only show an effect in activated fibroblasts. We found that when activated fibroblasts lack SEMA3C the fibrotic process, although it is not abolished, it is significantly reduced.

In an attempt to uncover the receptor involved in SEMA3C actions we first evaluated the expression of the different receptors in liver fibrosis and more specifically in HSCs. We found that only NRP2 was still expressed in activated HSCs. In keeping with this, deletion of NRP2 strongly inhibited HSC activation. Moreover, the deletion of NRP2 exclusively in myofibroblasts was sufficient to reduce fibrosis progression in mice, suggesting that activated HSCs respond to SEMA3C overexpression through NRP2 to mediate liver fibrosis. This discovery opens a new line of research, aiming at NRP2 blockade, which, together with the previously established benefits of NRP1 blockade(19,20), could serve as a therapeutic strategy to reduce liver fibrosis. Taken together, we demonstrate that SEMA3C exacerbates liver fibrosis. Its expression in HSCs is not only a marker of their activation, comparable to other previously defined, but also a mediator of this activation. Indeed, we demonstrate that HSCs decrease their level of activation upon SEMA3C deletion, opening the possibility to target this secreted protein to diminish liver fibrosis in patients.

## Abbreviations

CLD: chronic liver disease
NASH: non-alcoholic steatohepatitis
HSCs: hepatic stellate cells
NRP: Neuropilin-1
SEMA: semaphorin
HBV: chronic viral hepatitis B; hepatitis C HCV
HFD: High-fat diet
FPC: Fructose-Palmitate-Cholesterol
CCl4: carbon tetrachloride; y
IVC: inferior vena cava
GSEA: Gene Set Enrichment Analysis
ES: enrichment plot
NES: normalized enrichment score
FDR: False Discovery rate
ECs: endothelial cells.

## Acknowledgements

A special thank is going to the animal care takers of the Central Animal Unit in DKFZ and the Light Microscopy unit of the DKFZ and in particular to Dr. Damir Krunic for his help in image analysis. This work was funded by the Deutsche Forschungsgemeinschaft (DFG) project number 394046768 - SFB1366 projects C4 and Z2 (to A.F., C.M.), DFG project number 419966437, Deutsche Krebshilfe project number 70113888 (to J.R.V.), Deutsche Krebshilfe project 70113888, MCIN/AEI/ 10.13039/501100011033 (PID2020-117946GB-I00 and RYC2019-027937-I) (to J.R.V.) and (RTI2018-094734-B-C21) (to WJ), by “ERDF A way of making Europe” program and by “ESF Investing in your future”. Part of the equipment used in this work has been funded by Generalitat Valenciana and co-financed with ERDF funds (OP ERDF of Comunitat Valenciana 2014-2020).

## Author contribution

FDR: Conceptualization, Formal analysis, Investigation, Methodology, Validation, Visualization, Writing – review & editing; LW: Conceptualization, Formal analysis, Investigation, Methodology, Validation, Visualization, Writing – review & editing; MH: Formal analysis, Investigation, Methodology, SSH: Investigation, Methodology; IM: Formal analysis, Investigation, Methodology; SC: Formal analysis, Investigation, Methodology, WJ: Resources, Funding acquisition; MS: Investigation, Methodology; AB: Resources, BM-S: Resources, GW Resources, Investigation, Methodology; BE-Ü: Resources, Investigation, Methodology; SH: Resources; CM: Funding acquisition, Formal analysis, Resources; AF: Conceptualization, Formal analysis, Funding acquisition, Methodology, Validation, Visualization, Writing – review & editing, JR-V: Conceptualization, Formal analysis, Investigation, Methodology, Funding acquisition, Validation, Visualization, Writing – original draft.

