## Supplemental data for "Semaphorin 3C exacerbates liver fibrosis"

**Supplementary Table 1. Antibody list**

| **Protein** | **Company** | **Reference** | |
| --- | --- | --- | --- |
| αSMA | SIGMA | | A5228 |
| *SEMA3C* | Bio-Techne | | AF1728 |
| *DESMIN* | Abcam | | ab15200 |
| *SM22α* | Abcam | | ab137453 |
| *NRP1* | R&D Systems | | AF566 |
| *NRP2* | R&D Systems | | AF567 |
| *VCP* | Abcam | | ab109240 |
| *FN1* | Abcam | | ab2413 |
| Donkey anti-mouse IgG Alexa Fluor 546 | Invitrogen | | A10036 |
| *rabbit anti-mouse* | DAKO | | P0260 |
| *goat anti-rabbit* | DAKO | | P0448 |
| *rabbit anti-mouse* | DAKO | | P0260 |
| *rabbit anti-goat* | R&D Systems | | HAF017 |

**Suppl. Table 2. Primer list**

| **Gene** | **Forward Primer** | **Reverse Primer** |
| --- | --- | --- |
| *Acta2* | GAGAAGCCCAGCCAGTCG | CTCTTGCTCTGGGCTTCA |
| *Col1a2* | GTAACTTCGTGCCTAGCAACA | CCTTTGTCAGAATACTGAGCAGC |
| *Cph* | ATGGTCAACCCCACCGTG | TTCTTGCTGTCTTTGGAACTTTGTC |
| *Ctgf* | CTTCTGCGATTTCGGCTCC | TACACCGACCCACCGAAGA |
| *Hprt* | TGACACTGGCAAAACAATGCA | GGTCCTTTTCACCAGCAAGCT |
| *Mmp12* | CTCATGATGATTGTGTTCTTACAGG | GACAAGTACCATTCAGCAAATTCAC |
| *Nrp1* | GTCTTCAGGGCCCTTTCTCT | ATGTAGGTGCACTCCAAGCT |
| *Nrp2* | GCTGGCTACATCACTTCCCC | CAATCCACTCACAGTTCTGGTG |
| *Pai-1* | TCGTGGAACTGCCCTACCAG | ATGTTGGTGAGGGCGGAGAG |
| *PlexinD1* | GACTCGAACCTTCTTCCCCA | CAGTGAGGAGAACAGGCTGA |
| *S100a6* | AAGCTGCAGGATGCTGAAAT | CCCTTGAGGGCTTCATTGTA |
| *Sema3C* | TGGCCACTCTTGCTCTAGGT | AGATGCCTGTGGAGACTTGG |
| *Tagln* (SM22α) | TCCAGTCCACAAACGACCAAGC | GAATTGAGCCACCTGTTCCATCT |

**Supplementary Table 3. Cirrhosis signature**

| Rank | Gene | Rank | Gene | Rank | Gene | Rank | Gene | Rank | Gene |
| --- | --- | --- | --- | --- | --- | --- | --- | --- | --- |
| 1 | STMN2 | 43 | HKDC1 | 85 | GPR34 | 127 | KCNJ15 | 169 | OAS2 |
| 2 | KRT23 | 44 | ITGA2 | 86 | TMIGD3 | 128 | SPIC | 170 | FNDC5 |
| 3 | GPC3 | 45 | GSTM5 | 87 | IGHV1OR15-1 | 129 | LPAR1 | 171 | ITGA8 |
| 4 | IGKV1D-27 | 46 | PRKAA2 | 88 | FBLN5 | 130 | IGKV1OR2-108 | 172 | RNU6-287P |
| 5 | IGKV2D-40 | 47 | CFAP221 | 89 | PLCXD3 | 131 | PMP22 | 173 | MIR145 |
| 6 | MOXD1 | 48 | CDH6 | 90 | MMP2 | 132 | EDIL3 | 174 | IGKV1OR-2 |
| 7 | UPP2 | 49 | MMP7 | 91 | EMP1 | 133 | POF1B | 175 | OLFML1 |
| 8 | EPCAM | 50 | IGKV1-12 | 92 | GFPT2 | 134 | IGKV1D-33 | 176 | ANXA3 |
| 9 | FABP4 | 51 | SLC51B | 93 | SLC22A15 | 135 | CRISP3 | 177 | HSPB8 |
| 10 | IGKV3D-11 | 52 | IGKV2-24 | 94 | CD24P4 | 136 | IFI6 | 178 | FCGR1A |
| 11 | F13A1 | 53 | IGHV4-28 | 95 | CLMP | 137 | FRZB | 179 | GABRP |
| 12 | SCN7A | 54 | AQP1 | 96 | SEMA3C | 138 | GXYLT2 | 180 | SRGAP1 |
| 13 | GSTT1 | 55 | NPNT | 97 | RNASE6 | 139 | ADGRG1 | 181 | LAIR1 |
| 14 | EFEMP1 | 56 | PLPP4 | 98 | MX1 | 140 | DPYSL3 | 182 | ITGB8 |
| 15 | ITGBL1 | 57 | DCDC2 | 99 | SPON1 | 141 | STC1 | 183 | 44623 |
| 16 | CCDC80 | 58 | IGHV4OR15-8 | 100 | ESRP1 | 142 | SLIT2 | 184 | IGHV3-15 |
| 17 | FAM3B | 59 | LIPH | 101 | AMPD1 | 143 | HLA-DQB1 | 185 | LYZ |
| 18 | BICC1 | 60 | SMOC2 | 102 | ANTXR1 | 144 | LAMA2 | 186 | PKHD1L1 |
| 19 | KCNJ16 | 61 | DTNA | 103 | ANKRD22 | 145 | PLP2 | 187 | LY86 |
| 20 | RNU6-162P | 62 | MFAP4 | 104 | MGP | 146 | DOK5 | 188 | C1orf198 |
| 21 | LOXL4 | 63 | KRT7 | 105 | CLDN11 | 147 | GLT8D2 | 189 | DNAH5 |
| 22 | ENPP5 | 64 | IGKV1-9 | 106 | LUM | 148 | SVBP | 190 | IGHV3-33 |
| 23 | IGKV3D-20 | 65 | GSTM1 | 107 | CPA3 | 149 | CD180 | 191 | SVEP1 |
| 24 | CCL18 | 66 | VCAN | 108 | TRAT1 | 150 | CXCR4 | 192 | SLC28A3 |
| 25 | MUC13 | 67 | SLC5A1 | 109 | SULT1C2 | 151 | LGALS3BP | 193 | IGKV2D-30 |
| 26 | IGKV6-21 | 68 | SLC1A3 | 110 | EPHA3 | 152 | GABRE | 194 | C12orf75 |
| 27 | CHST4 | 69 | PRR15L | 111 | MAMDC2 | 153 | ADGRG2 | 195 | PARM1 |
| 28 | CCL21 | 70 | THY1 | 112 | CHI3L1 | 154 | ACSM1 | 196 | IGKV1-13 |
| 29 | CFTR | 71 | TACSTD2 | 113 | SRPX2 | 155 | RAB25 | 197 | PHLDB1 |
| 30 | TMPRSS3 | 72 | NRXN3 | 114 | CHIT1 | 156 | PTGDS | 198 | PDGFRA |
| 31 | CCL19 | 73 | CCL20 | 115 | CXCL11 | 157 | H2AFY2 | 199 | MTHFD2 |
| 32 | IGKV2D-26 | 74 | ELOVL7 | 116 | SLAMF8 | 158 | CDH11 | 200 | LAMB1 |
| 33 | UBD | 75 | LCN2 | 117 | GAS2L3 | 159 | HSPB6 | 201 | SLFN13 |
| 34 | MMRN1 | 76 | IGLV6-57 | 118 | IL7R | 160 | PODN | 202 | PFKP |
| 35 | ENPP2 | 77 | MAP1B | 119 | TAGLN | 161 | AEBP1 | 203 | C3AR1 |
| 36 | EHF | 78 | TMC4 | 120 | THBS2 | 162 | FAP | 204 | F3 |
| 37 | METTL24 | 79 | PROM1 | 121 | CHRDL1 | 163 | AC136428.1 | 205 | JCHAIN |
| 38 | VWF | 80 | SLCO2A1 | 122 | SORT1 | 164 | CRYAB | 206 | EGLN3 |
| 39 | RGS1 | 81 | SNAI2 | 123 | GPNMB | 165 | NEXN | 207 | MMP9 |
| 40 | DPT | 82 | LRRC1 | 124 | FGF23 | 166 | VEPH1 | 208 | LRRC7 |
| 41 | LGALS3 | 83 | VTCN1 | 125 | HTATSF1P2 | 167 | RNY3P11 | 209 | PLAT |
| 42 | SPP1 | 84 | SLC44A3 | 126 | SAA1 | 168 | UNC93A | 210 | NALCN |
| Rank | Gene | Rank | Gene | Rank | Gene | Rank | Gene | Rank | Gene |
| 211 | LEF1 | 254 | PCSK5 | 297 | IGHV3-47 | 340 | TNFRSF21 | 383 | ATP1B3 |
| 212 | FMN1 | 255 | SNAP25 | 298 | TMOD2 | 341 | CD34 | 384 | GSTP1 |
| 213 | ADAM28 | 256 | AC034105.1 | 299 | SLC12A2 | 342 | CSGALNACT1 | 385 | CPVL |
| 214 | RERG | 257 | HILPDA | 300 | TIGAR | 343 | PACS1 | 386 | PLA2G2A |
| 215 | IGHV3-13 | 258 | FMO1 | 301 | CD84 | 344 | RASSF4 | 387 | COL1A2 |
| 216 | SPARCL1 | 259 | SHISA3 | 302 | NAV3 | 345 | CACNA1C | 388 | TIMD4 |
| 217 | RN7SL751P | 260 | ITGA6 | 303 | SLC7A6 | 346 | GPRC5B | 389 | PKM |
| 218 | TAAR3P | 261 | IFI27 | 304 | RASL11B | 347 | GLIPR1 | 390 | FCGR2A |
| 219 | OMD | 262 | FMOD | 305 | IKZF3 | 348 | SEMA3A | 391 | SPINT2 |
| 220 | SERPINA12 | 263 | FPR3 | 306 | B3GNT5 | 349 | MACC1 | 392 | ARRDC2 |
| 221 | RAB3B | 264 | CCDC3 | 307 | CD69 | 350 | ADA2 | 393 | TESC |
| 222 | PLCE1 | 265 | FIBIN | 308 | TSPAN15 | 351 | IGKV4-1 | 394 | CDC6 |
| 223 | IGKV1D-16 | 266 | RNU6-1039P | 309 | GPR183 | 352 | IKZF1 | 395 | COL6A3 |
| 224 | SRPX | 267 | IGHV3-7 | 310 | MYOF | 353 | TMEM154 | 396 | SEMA3D |
| 225 | CASTOR2 | 268 | BACE2 | 311 | HIST1H2BM | 354 | GAPT | 397 | CD1E |
| 226 | CARD6 | 269 | FAM169A | 312 | TLR2 | 355 | MITF | 398 | NFASC |
| 227 | PDGFD | 270 | MINDY4 | 313 | IGKV1D-8 | 356 | IGKV2-18 | 399 | PALLD |
| 228 | HLA-DQA1 | 271 | TMEM45B | 314 | CACNA2D1 | 357 | LTBP2 | 400 | RASSF3 |
| 229 | HLA-DRB5 | 272 | PDE1A | 315 | SLC34A2 | 358 | DIRAS2 | 401 | PAQR5 |
| 230 | HLA-DQA2 | 273 | ABCC4 | 316 | WASF3 | 359 | CDKN2AIPNLP2 | 402 | LGALS9 |
| 231 | FBN1 | 274 | TMEM156 | 317 | GSTM3 | 360 | SGIP1 | 403 | PDE3A |
| 232 | ADCY3 | 275 | CCR7 | 318 | SNORD115-23 | 361 | HIST1H4H | 404 | AKR1B10 |
| 233 | TNFSF8 | 276 | PRDM1 | 319 | SPINT1 | 362 | C1orf210 | 405 | FCMR |
| 234 | ST14 | 277 | GPR171 | 320 | FBLN1 | 363 | RNY1P14 | 406 | PCDHB15 |
| 235 | CLDN10 | 278 | FBXO32 | 321 | LDOC1 | 364 | PDZK1IP1 | 407 | SOD3 |
| 236 | VGLL3 | 279 | GPC4 | 322 | CCL28 | 365 | HIST1H2AG | 408 | KPNA2 |
| 237 | LXN | 280 | SLPI | 323 | EPO | 366 | HAND2-AS1 | 409 | CMKLR1 |
| 238 | FSTL3 | 281 | CDC42EP3 | 324 | MUC6 | 367 | PIGR | 410 | PPDPF |
| 239 | VLDLR | 282 | FOLR2 | 325 | CD200 | 368 | MFGE8 | 411 | CAPG |
| 240 | APH1B | 283 | SLC7A11 | 326 | SULF1 | 369 | TLR7 | 412 | MTND4LP22 |
| 241 | CYP1B1 | 284 | KIT | 327 | ZDHHC13 | 370 | EPB41L4A | 413 | LPL |
| 242 | MIR4435-2HG | 285 | GOLM1 | 328 | PTPN14 | 371 | ADAMTS3 | 414 | IRAK3 |
| 243 | WFDC2 | 286 | MCUB | 329 | NCR3LG1 | 372 | PAPPA2 | 415 | GPR83 |
| 244 | CLSTN2 | 287 | PTGIS | 330 | GRAMD1B | 373 | CTSE | 416 | PTPN13 |
| 245 | SLAMF6 | 288 | HEYL | 331 | C7 | 374 | INMT | 417 | GALNT15 |
| 246 | OGN | 289 | CDH13 | 332 | PIK3IP1 | 375 | HSPA2 | 418 | RTP4 |
| 247 | CAPN6 | 290 | MCTP1 | 333 | CCDC102B | 376 | COL4A1 | 419 | ENTPD1 |
| 248 | RGS9 | 291 | MAML2 | 334 | CD52 | 377 | NTS | 420 | CYTIP |
| 249 | PSORS1C3 | 292 | IGKV1D-42 | 335 | IGHM | 378 | TNFAIP8 | 421 | CLIP4 |
| 250 | TLR5 | 293 | ISLR | 336 | MNS1 | 379 | GGT5 | 422 | FCER1G |
| 251 | UCA1 | 294 | BMP5 | 337 | GCNT4 | 380 | TCTN2 | 423 | SLC35G3 |
| 252 | ANOS1 | 295 | PRELP | 338 | SLIT3 | 381 | S100A11 | 424 | HLA-DPA1 |
| 253 | MAP2 | 296 | CCR2 | 339 | HLA-DRB3 | 382 | CA12 | 425 | UBE2L6 |
| Rank | Gene | Rank | Gene | Rank | Gene | Rank | Gene | Rank | Gene |
| 426 | FAT1 | 441 | CYBRD1 | 456 | LAMA4 | 471 | NFE2L3 | 486 | P2RY13 |
| 427 | PLVAP | 442 | IGF2BP2 | 457 | ABI3BP | 472 | GSN | 487 | PRKG1 |
| 428 | ZNF382 | 443 | SERPINE2 | 458 | HIST1H3I | 473 | SULT1C4 | 488 | TAX1BP3 |
| 429 | NEDD4L | 444 | HAMP | 459 | AP002956.1 | 474 | ANXA2 | 489 | B3GNT3 |
| 430 | ANXA1 | 445 | LHFPL6 | 460 | PRTFDC1 | 475 | GULP1 | 490 | DGKH |
| 431 | OSBPL3 | 446 | OSMR | 461 | FCER1A | 476 | TTN | 491 | HK1 |
| 432 | CXCL10 | 447 | ADAMDEC1 | 462 | RBPMS | 477 | IGLV7-46 | 492 | ISYNA1 |
| 433 | DKK3 | 448 | IGHV3OR15-7 | 463 | MAPK8IP1 | 478 | DCK | 493 | DAGLA |
| 434 | COL1A1 | 449 | PTP4A3 | 464 | CARMIL1 | 479 | GJA1 | 494 | CSF1R |
| 435 | YBX3 | 450 | MX2 | 465 | SAA2 | 480 | VIM | 495 | CD27 |
| 436 | SEZ6L2 | 451 | LOX | 466 | TNFSF13B | 481 | APOL3 | 496 | TUSC3 |
| 437 | CNKSR1 | 452 | ESCO2 | 467 | APOBEC3C | 482 | RUNX1T1 | 497 | CCND2 |
| 438 | PODXL | 453 | ZNF844 | 468 | MUC3A | 483 | MILR1 | 498 | SEL1L3 |
| 439 | FLNA | 454 | FAM83B | 469 | RAP1GAP | 484 | CD74 | 499 | HEPH |
| 440 | IL1RL1 | 455 | CRISPLD2 | 470 | ADAMTS9 | 485 | DCLK1 | 500 | CD3G |

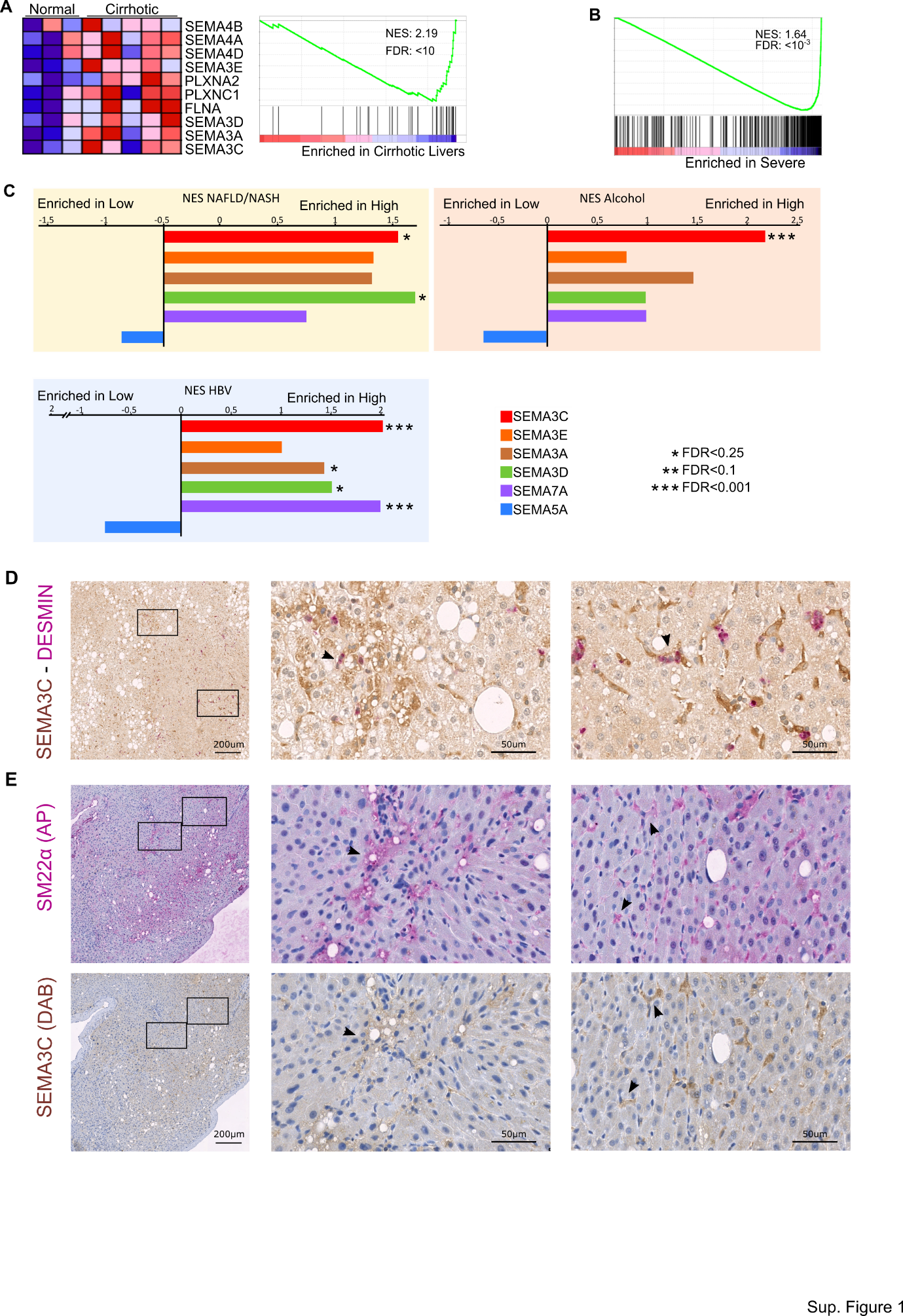

**Supplementary Figure 1. Gene set enrichment analysis (GSEA) of human liver samples. A** GSEA of the dataset GSE45050 using GO_SEMAPHORIN_PLEXIN_SIGNALING _PATHWAY gene set. Left panel shows a heatmap of the 10 genes most differentially enriched with the most enriched gene at the bottom. Right panel shows the enrichment plot including the Normalization Enrichment score (NES) and false discovery rate (FDR). **B** GSEA of the datasets GSE49541 using “Cirrhosis signature” gene set obtained A. Patients were classified according their fibrosis grade as mild (0/1) or severe (3/4). Left panel shows a heatmap of the 10 genes most differentially enriched with the most enriched gene at the bottom. Right panel shows the enrichment plot including the Normalization Enrichment score (NES) and false discovery rate (FDR). **C** GSEA of the datasets GSE61260 GSE103580 GSE83898 using “Cirrhosis signature” gene set obtained from Figure 1A. Patients were classified according their expression levels of SEMA3C, SEMA3A, SEMA3E, SEMA3D, SEMA7A or SEMA5A as over (high) or under (low) the median expression of each gene. Graphs represent FDR values. * FDR<0.25, ** FDR<0.1, *** FDR<0.001. **B** SEMA3C expression in the different cell types composing the liver, comparing uninjured (grey) with cirrhotic (purple) donors. **D** Co-immunohistochemistry for DESMIN (stained by alkaline phosphatase AP) and SEMA3C (stained by DAB) were performed on biopsies of human livers. Samples were counterstained with hematoxylin. Both proteins were closely located (arrows). **E** Staining of SEMA3C (DAB) and SM22α (AP) on serial sections from biopsies of human livers. Samples were counterstained with hematoxylin. Arrows show representative areas with positive staining. Scale bars are 200μm in the zoom-out images and 50μm in the zoom-in images.

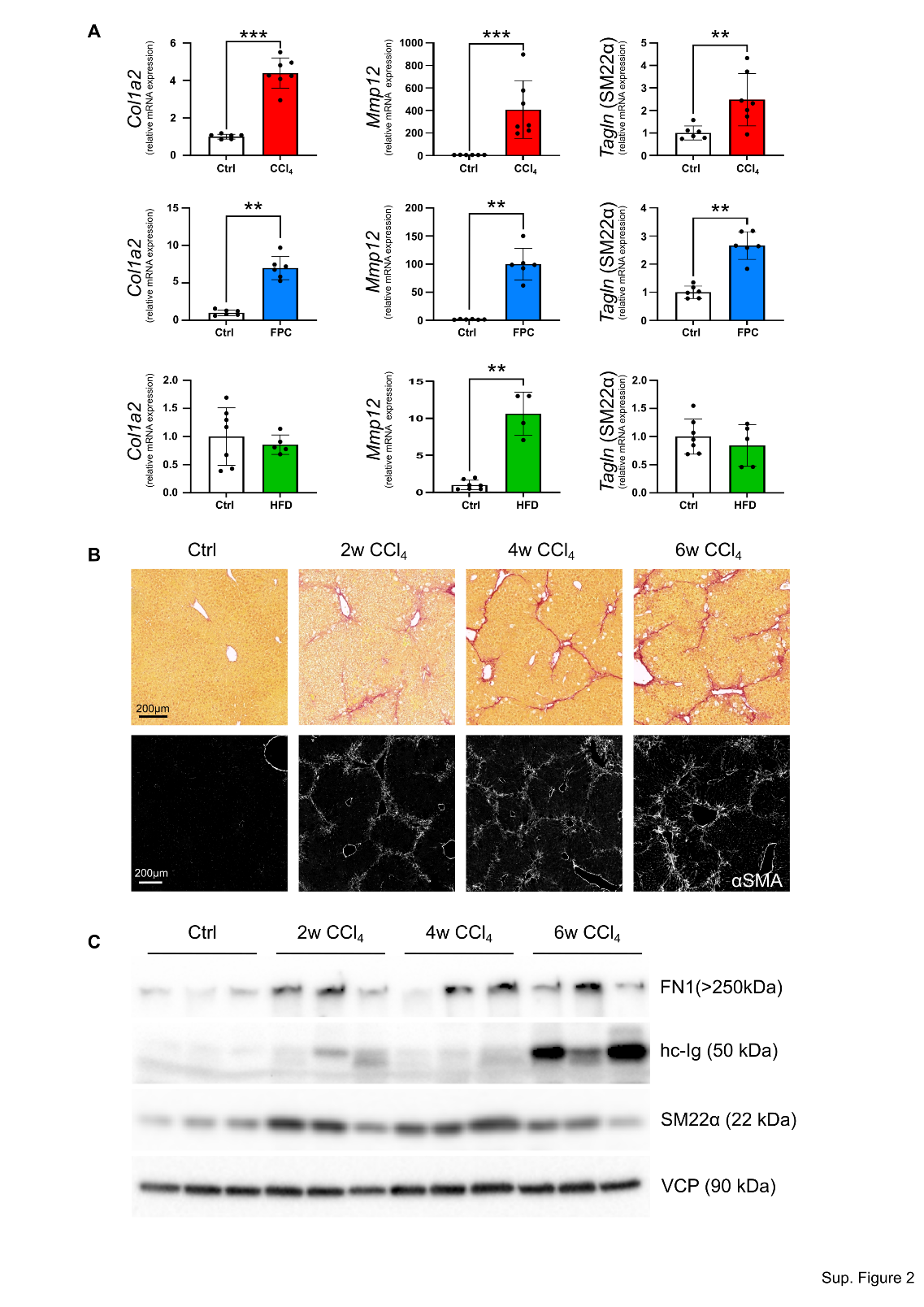

**Supplementary Figure 2. Characterization of liver fibrosis.** **A** The gene expression of liver fibrosis markers (*Col1a2*, *Mmp12* and *Tagln*) was analysed by qRT-PCR. CCl_4_, FPC and HFD data (respectively red, blue and green columns) are shown as relative to their own controls (white columns). One-tailed non-parametric Mann-Whitney test was performed to evaluate data significance. * and ** correspond to p value ≤0.05 and ≤0.01, respectively. To determine the progression of liver fibrosis and to establish the timing for CCl_4_ treatment, **B** Sirius red and alpha-smooth muscle actin (αSMA) staining was performed on livers treated with CCl_4_ for 2, 4 or 6 weeks and compared to non-treated mice. Scale bars are 200 µm. The same samples were also analysed by Western blot for fibronectin (FN1), heavy-chain immunoglobulin (hc-Ig), SM22α and the loading control Valosin-containing protein (VCP) (**C**).

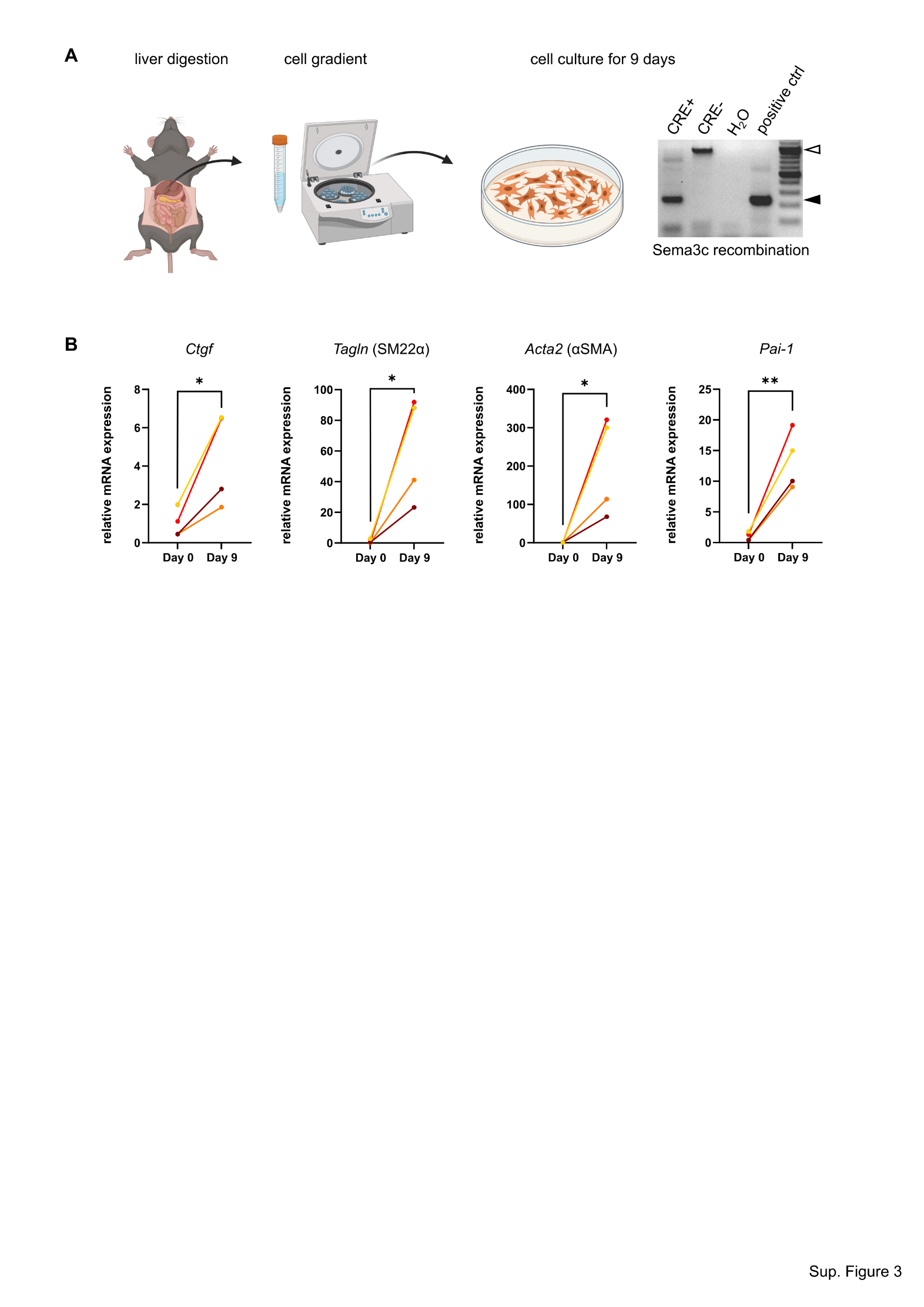

**Supplementary Figure 3. HSC activation.** **A** Scheme of the workflow for HSC isolation. After liver digestion, HSC are isolated by cell gradient and cultured for 9 days. Before to be used in experiments, HSC cells are tested for gene recombination by PCR. White arrow indicates the band that is obtained in absence of recombination (CRE-, control mice); while black arrow indicates gene recombination and the consecutive Sema3C knock-out. **B** Gene expression of activation markers *Ctgf*, *Tagln* (SM22α), *Acta2* (αSMA) and *Pai-1* was analysed in WT-HSCs upon cell culture. One-tailed paired t-test was performed to evaluate data significance. * and ** correspond to p value <0.05 and <0.01, respectively.
